# Wolbachia has subtle effects on thermal preference in highly inbred Drosophila melanogaster which vary with life stage and environmental conditions

**DOI:** 10.1101/2023.02.17.528917

**Authors:** Anton Strunov, Charlotte Schoenherr, Martin Kapun

## Abstract

Temperature fluctuations are challenging for ectotherms which are not able to regulate body temperature by physiological means and thus have to adjust their thermal environment via behavior. However, little is yet known about whether microbial symbionts influence thermal preference (*T*_p_) in ectotherms by modulating their physiology. Several recent studies have demonstrated substantial effects of *Wolbachia* infections on host *T*_p_ in different *Drosophila* species. These data indicate that the direction and strength of thermal preference variation is strongly dependent on host and symbiont genotypes and highly variable among studies. By employing highly controlled experiments, we investigated the impact of several environmental factors including humidity, food quality, light exposure, and experimental setup that may influence *T*_p_ measurements in adult *Drosophila melanogaster* flies. Additionally, we assessed the effects of *Wolbachia* infection on *T*_p_ of *Drosophila* at different developmental stages, which has not been done before. We find only subtle effects of *Wolbachia* on host *T*_p_ which are strongly affected by experimental variation in adult, but not during juvenile life stages. Our in-depth analyses show that environmental variation has a substantial influence on *T*_p_ which demonstrates the necessity of careful experimental design and cautious interpretations of *T*_p_ measurements together with a thorough description of the methods and equipment used to conduct behavioral studies.

## Introduction

Temperature modulates many physiological processes and has direct effects on development, survival and reproduction of any organism^1, 2^. Ectotherms lack the ability to regulate their body temperature by physiological means and are particularly affected by temperature variation^3^. Their thermoregulation is thus often mediated through behavior^4^ (Stevenson, 1985) so ectotherms tend to occupy environmental niches close to their optimal thermal conditions to survive and propagate. Every organism exhibits a thermal preference (*T_p_*), which is the preferred body temperature or temperature range that is chosen in the absence of other ecological constraints^5^. However, estimates of *T*_p_ are by no means absolute, but may be strongly influenced by interactions with ecological factors and even with symbionts. Endotherms, for example, combat bacterial and viral infections with fever by physiologically increasing their body temperature outside the thermal optimum of their infectious agents. Several studies show that also ectotherms employ similar behavioral strategies by changing their *T*_p_^6–11^. For example, crickets actively seek higher temperatures when infected with the pathogenic bacterium *Serratia marcescens*, which may have physiological effects similar to fever in response to pathogenic infections^12^. However, also the opposite behavior, i.e., behavioral chill, has been reported for poikilotherm organism as a mechanism to fight pathogens, for example, in *Drosophila*^13^.

In particular, three recent studies investigated changes of thermal behavior in *Drosophila melanogaster* infected with the bacterial endosymbiont *Wolbachia*^14–16^. Two of these studies found that, depending on the *Wolbachia* variant, infected flies chose colder temperatures than uninfected individuals^14, 15^. While such a behavioral response is probably strongly influenced by environmental and experimental conditions and not a general pattern^16^, these findings may suggest that the hosts can alleviate detrimental effects of high titer infections by choosing ambient temperatures outside the optimal physiological range of the symbiont. It, however, remains uncertain if such behavioral patterns are restricted to adults or are also found at juvenile life stages. Finding the optimal temperature for development is pivotal since any major disturbance during juvenile stages might severely decrease survival and reproduction chances of the imago^17^. Manipulating thermal preference at this life stage of the host might thus be risky yet beneficial for the microbial symbiont, by “setting up” its thermal environment early in the host development. However, to the best of our knowledge, no study has yet investigated the effects of *Wolbachia* on *Drosophila T*_p_ at juvenile life stages.

Measuring thermal behaviour in *Drosophila* is complex, and many environmental factors and experimental noise that potentially influence the *T*_p_ have to be accounted for. These can include, for example, environmental factors such as humidity, light, olfactory and acoustic cues, but also properties of the experimental design, such as age of the flies, developmental temperature, number of specimens and the design of the apparatus that measures thermal preference, as comprehensively reviewed in Dillon et al.^18^. This issue becomes evident when comparing the divergent results of the three aforementioned studies that specifically assessed the *T*_p_ of *D. melanogaster* in the presence of *Wolbachia* infections^14–16^. The absolute *T*_p_ values for the same *Wolbachia* strain vary strongly among the studies (up to 5-6°C), which is potentially influenced by different host genetic backgrounds, experimental conditions or temperature ranges used in these studies. Assessing the effects of different environmental factors on thermal preference in the *Wolbachia*-*Drosophila* system is therefore crucial for an informed analysis and interpretation of behavioral measurements.

Here, we demonstrate that several environmental factors, such as humidity, food, and the structure of the thermal gradient device have a major impact on *T*_p_ in adult *D. melanogaster* flies in the context of strain-specific *Wolbachia* infections. Additionally, we investigate - for the first time - the effect of *Wolbachia* on juvenile *T*_p_ and find that it is not affected by infection in early and late fly larvae and at pupariation.

## Materials and Methods

### Fly lines

In our experiments, we used four highly inbred, long-term laboratory strains of *D. melanogaster* which were uninfected (*w*-) or infected with either two of the most common natural *Wolbachia* variants (*w*Mel and *w*MelCS) or with the *w*MelPop lab-variant that were previously investigated by Truitt et al.^14^ for thermal preference. All fly lines were initially established by Luis Teixeira^19^ in a DrosDel *w*^1118^ isogenic background. Flies were maintained on a custom fly medium based on agar-agar, molasses, and yeast at 24°C with 12h:12h light:dark cycle and an average humidity of 50%. Prior to experiments, the infection status was confirmed by PCR using *Wolbachia*-specific primers amplifying parts of the *wsp* gene: forward - tggtccaataagtgatgaagaaactagcta and reverse - aaaaattaaacgctactccagcttctgcac^20^. *Wolbachia* variants infecting the fly strains were distinguished with diagnostic VNTR-141 PCR primers: forward - ggagtattattgatatgcg and reverse - gactaaaggttattgcat. PCR products vary in length among *Wolbachia* strains^21^.

### Thermal gradient machine

To carry out *T*_p_ measurements in *D. melanogaster* larvae, during pupariation and in adult flies, we designed and built a new thermal gradient apparatus with two narrow arenas for precise temperature measurements and with limited space for movement along the thermal gradient. The “plate” device consisted of an elongated aluminum plate as the bottom piece resting on two peltier elements, which are either heating or cooling to generate a temperature gradient. On top, we used an equisized plexiglass plate as the cover, which contained holes to apply test subjects. Two narrow arenas with 6 mm heights and widths were established by placing three thin plexiglass dividers between the two plates, which contained holes for inserting temperature sensors. We restricted the length of the arena to 30 cm by adding foam stoppers at the ends of each lane to prevent flies from escaping and to maintain airflow. In addition, Whatman paper was used as a bottom layer to cover the aluminum plate in the arenas. We built two identical devices with the aforementioned design, which allowed us to investigate *T*_p_ in four arenas in parallel during one experimental assay (see https://github.com/capoony/DrosophilaThermalGradient for a detailed description).

For *T*_p_ measurements in adult flies, we additionally designed and built a second “tube” thermal gradient device with a tubular shape following Rajpurohit and Schmidt^22^ and complementing the design as described in Truitt et al.^14^. This apparatus consisted of a central cylindrical aluminum rod connected to quadratic aluminum bases at both ends that conduct heat from two peltier elements, which are either heating or cooling. The aluminum rod was placed inside a transparent plexiglass tube, which left an open space of approximately 4 cm in diameter. The plexiglass tube was attached to the aluminum rod with Teflon covers which enclosed the fly arena on both sides. The plexiglass tube contained several holes to insert flies and to attach sensors for temperature measurement. A Whatman paper was inserted right above the aluminum rod to divide the arena into two halves. Flies were kept in the upper half, which facilitated imaging the location of the flies within the cylindrical arena around the central rod from atop during *T*_p_ measurements (see https://github.com/capoony/DrosophilaThermalGradient for a detailed description).

Peltier elements (P&N Technology, Xiamen Fujian China) generating heat were attached to small heat sinks (Fischer Electronic, Germany) connected to a ventilator (Oezpolat, Germany) to stabilize the temperature. Cooling peltier elements produce excess waste heat and were therefore placed on top of a bigger heat sink (Fischer Electronic, Germany), which was partially submerged in a bowl with cold water that was constantly cooled down by pipes supplying cold water from a distant water bath (fbc630, Fisher scientific, USA).

20 min prior to each experiment, we attached the peltier elements to power supplies set to 2 V and 3 V for cooling and heating, respectively, to stabilize the temperature gradient within the thermal gradient apparatus. We found that the temperature gradient remained stable for at least 1 h, which generally exceeded the duration of an experimental run (app. 20-30 min). After each experiment, the plexiglass parts of the apparatus were cleaned with soap and hot water to remove any potential pheromones and other odorants, which might interfere with thermal preference behavior of flies.

Each experiment was carried out in a dark isolated room with no, or very low noise distraction. The room was equipped with air-conditioning to keep the ambient temperature stable at 21-23°C, which was constantly monitored with a data logger. The temperature within the devices was measured at three points (in the center and at the hot and cold edges) with digital thermal sensors (Analog devices, USA). The data from thermal sensors were collected with a custom python script on a Raspberry pi 3B+ computer (Raspberry Pi Foundation, UK) in 10 sec intervals and saved as text files. The position of the flies within the devices were assessed by an infrared camera attached atop of the thermal gradient apparatus. Images were taken every 30 sec and saved to the Raspberry pi computer for each run. We developed a custom python script to estimate *T*_p_ for every fly using information of an individual’s position relative to the linear temperature gradient between the temperature measurement points. The coordinates of each individual fly and of the thermal sensors were manually assessed in ImageJ^23^ based on infrared images captured 20 min after the onset of each experiment. The python scripts for quantifying thermal preference from the coordinates including a test dataset, a detailed description and all raw images from the thermal preference assays are available at GitHub (https://github.com/capoony/DrosophilaThermalGradient).

### Temperature preference measurements in larvae

To assess larval *T*_p_, we used early and late 3^rd^ instar larvae, 72 h, and 120 h after egg laying (AEL), respectively, and focused on two fly strains that were either uninfected (w-) or infected with *w*MelCS for thermal preference experiments (see *Fly lines* section in Materials and methods). To obtain tightly synchronized cohorts of larvae, we maintained 40-50 adult female flies at 24°C in glass vials with the medium for 3-6 h for egg laying and then transferred these to a fresh medium for collecting another replicate. After 72 or 120 h, larvae were moved to an 18% sucrose solution (in dH_2_O) using a spatula. Floating larvae were gently collected with a cut 1 mL pipette tip and transferred to dH_2_O. After washing three times with dH_2_O, larvae were transferred into empty plastic Petri dishes (d=2.5 cm) and kept for 10-20 min to recover from the stress.

For *T*_p_ measurements, we covered the bottom of each lane in the thermal gradient machine with a 1mm layer of 2% agarose gel to create a habitable environment for larvae that allowed them to crawl freely. We started the experiment when the thermal gradient was constant and linear from 18°C to 28°C (1°C per 3cm). Experiments were carried out in a dark room at 24-25°C, with 30% humidity at different times of the day (from 11 am to 4 pm). Each run included two replicates of the *w*- and the *w*MelCS lines. 50-100 larvae were introduced in the middle of each lane (23°C) with a wet brush and a Plexiglass cover was applied on top of the machine to prevent larvae from escaping. The position of larvae was analyzed on site after 30 min by quantifying the numbers of larvae located in a certain temperature zone marked with a pen beforehand.

### Measurement of pupariation Tp

For assessing *T*_p_ at pupariation, we used late 3^rd^-instar larvae, 140h after egg laying (AEL) close to pupariation. Again, we focused on the *w*- and *w*MelCS strains for the experiments and obtained synchronized cohorts of 140 h-old larvae as explained above.

For *T*_p_ measurements, we covered the bottom of the four lanes of the gradient machine with a 1 mm thick sheet of odorless black plastic to create a contrasting background and to supply a solid surface for pupariation. We inserted larvae when the gradient was constant and linear from 18°C to 28°C (1°C per 3cm). Experiments were carried out in a dark room at 24-25°C with 30% humidity for approximately 12h until every larva had pupariated. Each run included two replicate lines at the same time (2 times *w*- and 2 times *w*MelCS). 50-100 larvae were introduced in the middle of the lane (23°C) by wet brush and a plexiglass cover was used to seal the top of the machine to prevent larvae from crawling out of the arenas. Thermal preference at pupariation was analyzed after 6-12 h by quantifying the numbers of pupae in certain temperature zones as described above.

### Temperature preference measurements in adults

Prior to the experiments, we selected cohorts of 20 females and 20 males, which were one week old (± two days), for each of the four strains (*w*-, *w*Mel, *w*MelCS and *w*MelPop). We transferred these to fresh glass vials with medium by gently applying CO_2_ and then allowed the flies to recover for two days. To reduce the possibility of an experimenter bias, we anonymized the ID’s of the cohorts prior to experiments by an independent researcher. For each assay on the “plate” device, we included one replicate of all four fly lines and placed each of the groups in one of the four lanes. For experiments using the “tube” device, we carried out two replicate assays in parallel, using randomly drawn cohorts. Flies were gently collected from the vial with a handmade fly aspirator and transferred to the thermal gradient apparatus through a hole in the center. Then, the hole was tightly sealed with cotton, the light was turned off and measurements were taken during 20 min. An image of the whole apparatus was taken every 30 seconds with an infrared camera attached atop the setup.

Because time of the day can have a strong influence on *T*_p_ in *Drosophila*^24^, we performed a maximum of 4 runs per day to avoid potential bias from circadian rhythms. The time of the runs during the day was usually as follows: first two runs from 10 am to 12 am, last two runs from 1 pm to 3.30 pm. To quantify *T*_p_ of individual flies in the arena, we used the images of fly locations taken 20 min after the onset of the experiment similar to Truitt et al.^14^.

In addition to the experiments outlined above, we sought to test how environmental variation influences *T*_p_. We accounted for four different factors: humidity, light, food, and type of the gradient machine. All four factors were tested in at least three replicates with the “tube” device, except for the last one where both types were used for comparison. When testing for the effects of humidity, we used a standard humidifier to increase the humidity from 30%, which we usually measured without any adjustment, to stable 60% humidity in the test room. To test the influence of light on thermal preference, we positioned the gradient machine at the center above a ceiling lamp and performed thermal preference runs with the light on and off using different biological replicates. When testing for the effect of food quality on *T*_p_, we kept the flies for at least three generations on different diets before conducting the experiment. We compared the effects of our in-house food recipe, described above, to Carolina 4-24 instant food, obtained directly from the manufacturer (Carolina, USA). To test how the type of device affects *T*_p_, we combined the data of the runs in the two different devices carried out in the same room with similar ambient temperatures to keep environmental conditions consistent. The analysis of images taken under infrared light did not allow to distinguish between sexes. Since differences in *T*_p_ has not been reported in *Drosophila melanogaster*^25, 26^, we do not assume that ignoring the factor “sex” would lead to biased results.

### Statistical analyses

The effects of *Wolbachia* infection and other environmental factors on *T*_p_ were analyzed using generalized linear mixed models (GLMMs) with a Poisson error structure in *R*^27^ based on the “glmer” function in the lme4 package^28^. In our models, we considered *T*_p_ of every individual as the dependent variable, and infection type as the fixed factor. Moreover, we always included the factor replicate experiments nested within infection type as random factors in each of our models. For the larval *T*_p_ experiment, we additionally included the larval age (72 h and 120 h) as a fixed factor and the interaction of larval age with infection status in our model. Moreover, for most of the models, we included the time of the run of each replicate during the day as a random factor. We used the *anova* function in *R* to test for a significant effect of infection status on thermal preference and calculated a Type-III analysis of deviance comparing nested models to test for significant effects of fixed factors and interaction in the *T*_p_ experiments. Whenever the factor *infection status* had a significant effect, we performed post-hoc tests using Tukey’s HSD method as implemented in the *emmeans* package^30^. We also inferred the skewness of *T*_p_ distributions using the *moments* package^31^ and compared the average skewness among experimental setups with Wilcoxon Rank tests in *R*. All *R* code can be found at https://github.com/capoony/DrosophilaThermalPreference.

## Results and Discussion

### Wolbachia infection has no effect on T_p_ in 3^rd^-instar Drosophila larvae and on pupariation T_p_ in D. melanogaster

Using the newly designed thermal gradient apparatus (see Materials and Methods), we found that *T*_p_ in early (72h AEL; n=281) and late (120h AEL; n=658) 3^rd^-instar larvae significantly shifted from a median *T*_p_ of 22-23°C to a median *T*_p_ of 19°C (see **Table S1**) with progression of development (GLMM: *p*=2.8e-8, *x*^2^=34.8, *df*=2; **Table 1A, Figure 1A**). Our results were consistent with previous findings^32, 33^. However, we neither observed an effect of *Wolbachia* on *T*_p_ in 3^rd^-instar larvae (GLMM; *p*=0.34, *x*^2^=2.1, *df*=2; **Table 1A; Figure 1A**) nor an interaction of *Wolbachia* with larval age (GLMM; *p*=0.17, *x*^2^=1.9, *df*=1). Since many larvae clustered at the coldest spot at the wall in the gradient machine (18-19°C), we repeated the experiment for the 120h age class, which exhibited lower *T*_p_, and extended the lower temperature range to 15°C. This extended analysis of 529 infected and 332 uninfected 3^rd^-instar larvae at 120h AEL similarly showed no effect of *Wolbachia* infection on *T*_p_ of the host (GLMM: *p*=0.38, *x*^2^=0.76, *df*=1; **Figure S1**, **Table 1B**), although the median *T*_p_ decreased by approximately 1°C with the majority of larvae of both infection types (*w*- and *w*MelCS) clustering at temperatures between 16°C and 18°C (median *T*_p_ of 18°C; **Table S1**).

**Figure 1.**
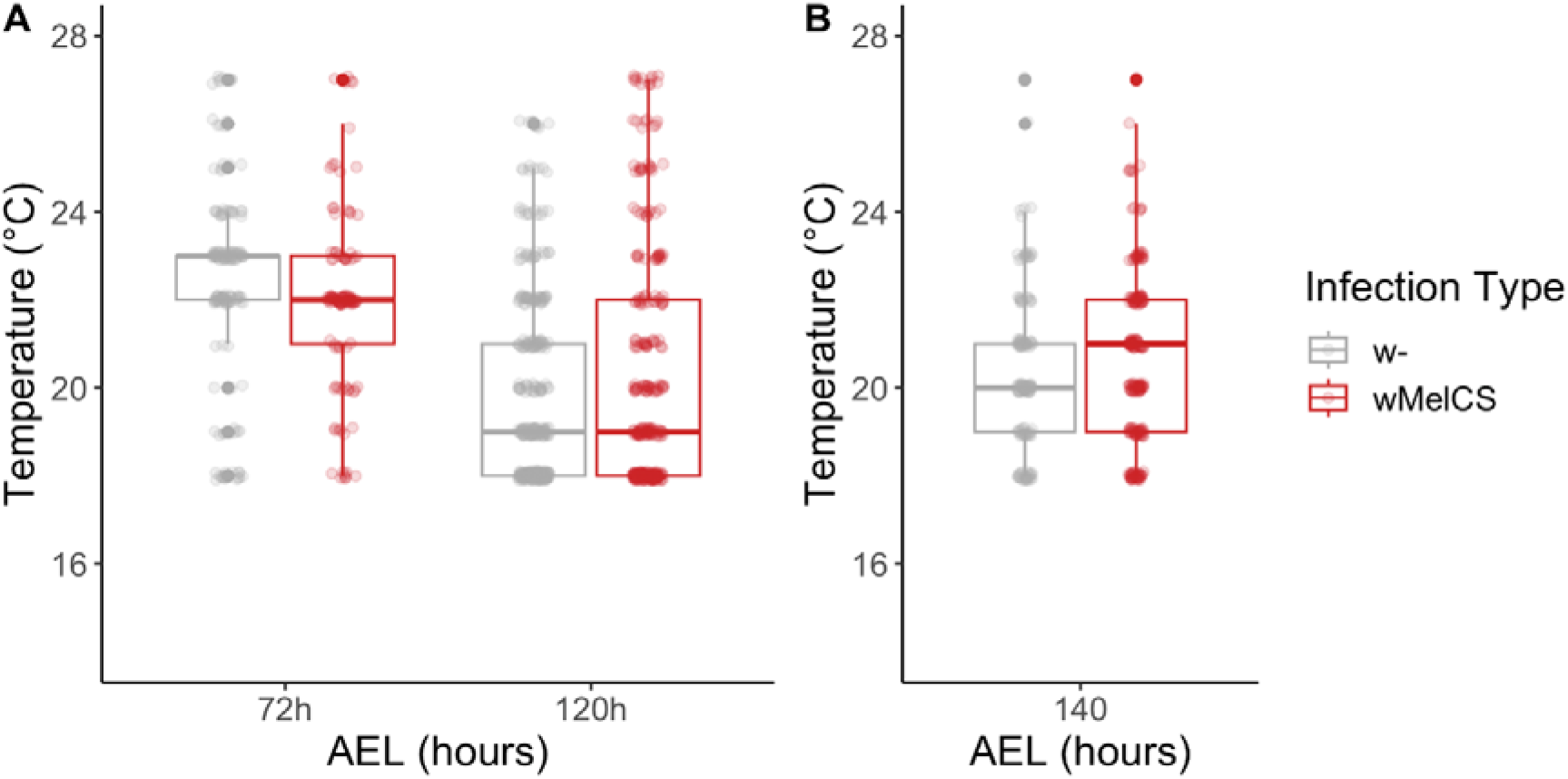
Thermal preference (*T*_p_) of early and late *Wolbachia*-infected *D. melanogaster* 3^rd^-instar larvae and larvae at the onset of pupariation. (A) Temperature shift in early and late 3^rd^-instar larvae that were either uninfected (w-; grey) or infected with wMelCS (red), 72 hours and 120 after egg laying (AEL) respectively (see also Figure S1 and Table 1). (B) *T*_p_ of *D. melanogaster* larvae at the onset of pupariation (140h AEL) for both infection types (see also Table 1). Note the absence of *Wolbachia* effects on thermal preference at all larval stages.

**Table 1.**
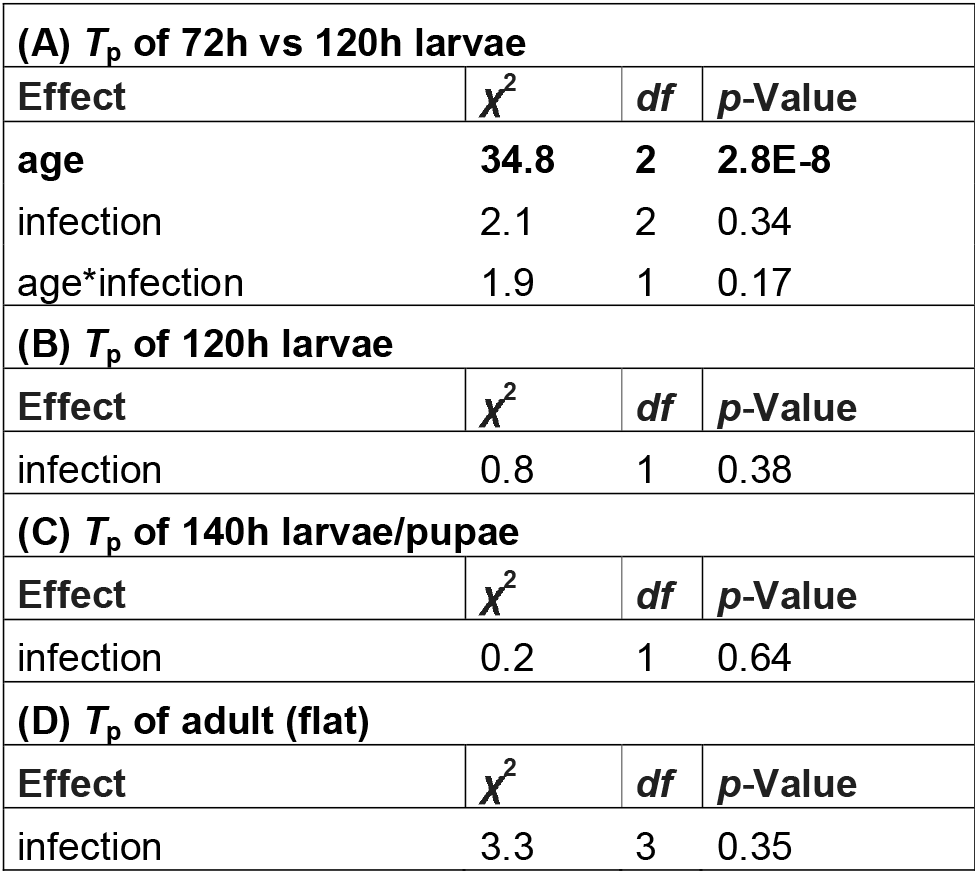
The influence of *Wolbachia* infections on *Drosophila T*_p_ at different stages of development. Significant effects of generalized mixed models (GLMM) inferred by analysis of deviance are highlighted in bold.

| **(A) $T_p$ of 72h vs 120h larvae** |  |  |  |
| --- | --- | --- | --- |
| Effect | $\chi^2$ | df | p-Value |
| **age** | **34.8** | **2** | **2.8E-8** |
| infection | 2.1 | 2 | 0.34 |
| age*infection | 1.9 | 1 | 0.17 |
| **(B) $T_p$ of 120h larvae** |  |  |  |
| Effect | $\chi^2$ | df | p-Value |
| infection | 0.8 | 1 | 0.38 |
| **(C) $T_p$ of 140h larvae/pupae** |  |  |  |
| Effect | $\chi^2$ | df | p-Value |
| infection | 0.2 | 1 | 0.64 |
| **(D) $T_p$ of adult (flat)** |  |  |  |
| Effect | $\chi^2$ | df | p-Value |
| infection | 3.3 | 3 | 0.35 |

Pupariation in *Drosophila* is a crucial developmental stage during which the fly is immobilized for several days and undergoes complete metamorphosis. Thus, for larvae to properly develop choosing the optimal pupariation temperature is of great importance^18, 34^. We analyzed the *T*_p_ of 192 infected and 142 uninfected larvae about to pupariate (140h AEL; **Table S1**). In comparison to *T*_p_ in late 3^rd^-instar larvae (median *T*_p_ of 18°C for both infection types; **Table S1**), the pupariation *T*_p_ shifted to warmer temperatures (**Figure 1B**, median *T*_p_ of 20°C for uninfected, and 21°C for infected individuals; **Table S1**). However, similarly to 3^rd^-instar larvae, we found no significant effect of *Wolbachia* on thermal preference at pupariation stage (GLMM: *p*=0.64, *x*^2^=0.21, *df*=1; **Table 1C**).

The absence of *Wolbachia* effects on *T*_p_ in early and late 3^rd^-instar larvae and on pupariation *T*_p_ might indicate that infection does not impose significant costs at least in the developmental stages investigated here. It might be correlated with bacterial titer, which is known to influence the fitness of symbiotic association when it reaches high levels^35–39^. Although little is known about *Wolbachia* titer dynamics during fly development and in spite of yet undiscovered effects of *Wolbachia* infections during larval development, our findings could imply dormancy of infection at juvenile stages of the host. Strunov et al.^37^, for example, showed that the titer of the pathogenic *w*MelPop strain in *D. melanogaster* is significantly lower in larvae than in the imago and stable across development without any effects of rearing temperature. However, an influence of temperature on *Wolbachia* titer has been previously observed in adults after metamorphosis^40, 41^, indicating that bacterial multiplication and pronounced interaction with the host may take place during late pupariation stage. In line with this hypothesis, it was shown that *Wolbachia* interferes with pheromone production in females during pupariation and affects female-to-male communication^42^. Additionally, two recent papers demonstrated the absence of *Wolbachia* effects on locomotion and fitness at juvenile stage of wild-caught *D. nigrosparsa*^43^ and *D. melanogaster* flies^44^. The absence of *Wolbachia* effects on host juvenile traits is challenged by a study, which reported stable abundance of *Wolbachia* transcripts at all stages of host development^45^, although there was no data on translation of the transcripts. Thus, *Wolbachia* have presumably no or very low and undetected effects on developing flies. Additionally, other cues like foraging for food^46^ might represent stronger behavioral stimuli, which could mask subtle effects of *Wolbachia* on thermal behavior.

### Wolbachia infection has subtle effects on thermal preference in adult flies that are influenced by the environment and experimental design

Using our new experimental setup, we sought to reproduce previous experiments which reported that *Wolbachia* infection can lower thermal preference in adult *D. melanogaster* flies depending on the *Wolbachia* variant studied^14, 15,but^ ^see^ ^16^. We used the same fly lines, initially established by Teixeira et al.^19^, as examined by Truitt et al.^14^. We investigated five replicate cohorts of each fly line (*w*-, *w*Mel, *w*MelCS, and *w*MelPop) with our new thermal gradient device under highly controlled environmental conditions. Unexpectedly, we were unable to reproduce the previous results in Truitt et al.^14^, since none of the lines differed in thermal preference (GLMM: *p*=0.35, *x*^2^=3.29, *df*=3; **Table1D**; **Figure 2**). Median *T*_p_ was 18.3°C, 18.8°C, 18.1°C, and 18.2°C for *w*-, *w*Mel, *w*MelCS, and *w*MelPop, respectively (**Table S1**). The variation among the replicates was quite high with 2.4 to 3°C standard deviation among the mean *T*_p_, depending on the infection type studied.

**Figure 2.**
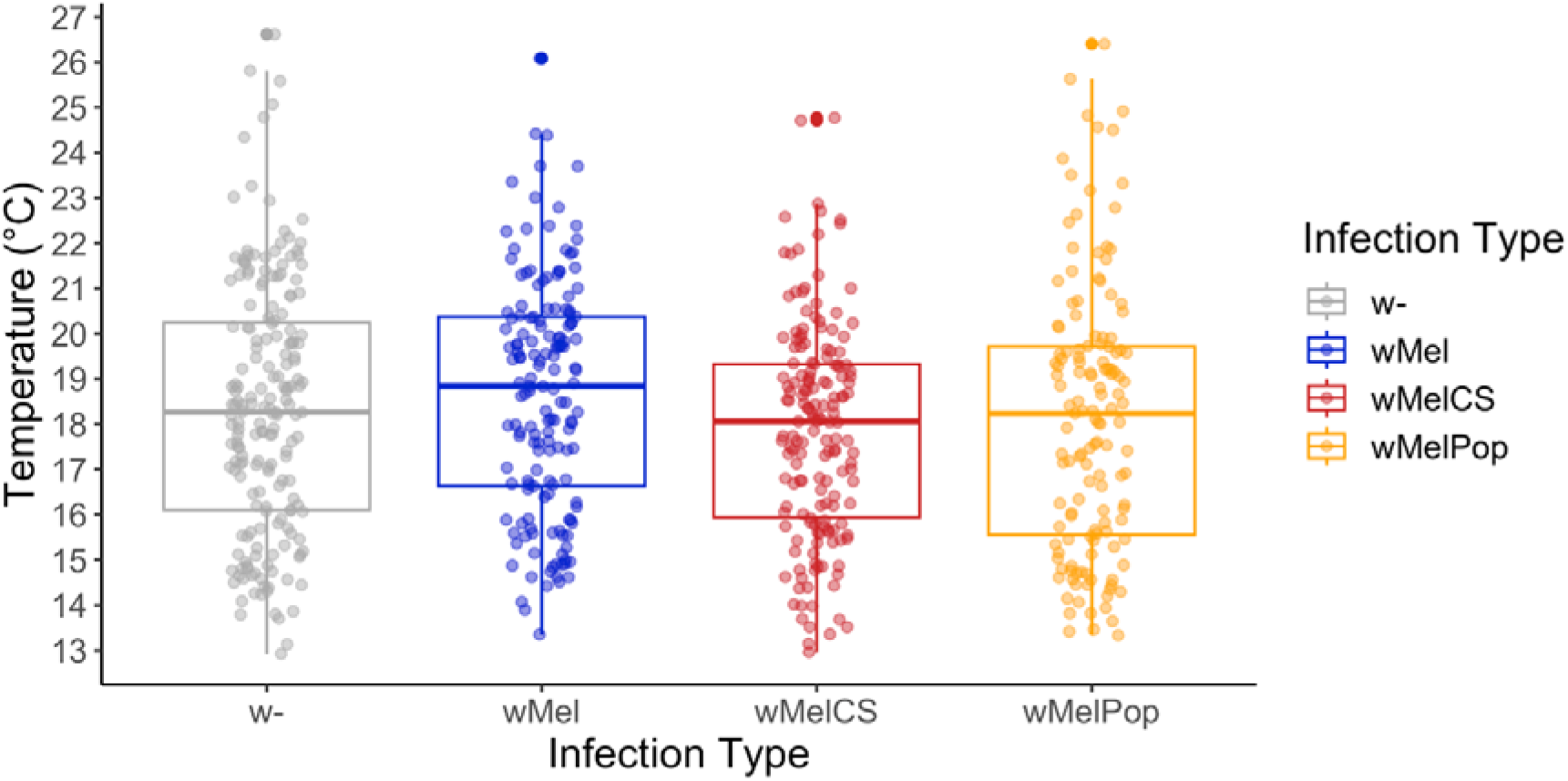
Thermal preference of adult *D. melanogaster* with different infection types. Boxplots showing the median *T*_p_ of flies that were either uninfected (w-) or with three different *Wolbachia* types (wMel: blue; wMelCS; red and wMelPop: orange)

To rule out that the absence of *T*_p_ variation among strains infected with different *Wolbachia* variants is an artifact of our experimental setup, we newly built a device similar to the design in Truitt et al.^14^ and repeated the *T*_p_ assays. Overall, median *T*_p_ was approximately 3°C higher than the measurements from the “plate” device (21.7°C, 22°C, 21.5°C and 21.5°C average *T*_p_ for *w*-, *w*Mel, *w*MelCS, and *w*MelPop, respectively; **Table S1**), which indicates a strong influence of the design on absolute *T*_p_ (GLMM: *p*<2.2e-16, *x*^2^=748.6, *df*=4; **Table 2A**). Additionally, we found significant differences among infection types (GLMM: p<0.0044, *x*^2^=18.88, *df*=3; **Table 2A**), albeit in the opposite direction according to our expectations: the *T*_p_ of the *w*Mel variant increased by approximately 1°C compared to *w*-(Tukey’s post hoc test: *p*<0.003; **Table S2**). In contrast, *w*MelCS and *w*MelPop variants did not show any effect on *T*_p_ compared to the uninfected control lines (Tukey’s post hoc test: *p*>0.05; **Table S2**), which is in stark contrast to substantial *T*_p_ differences (2-4°C) among variants previously described by Truitt *et al*.^14^. Thus, despite using a similar design for the thermal gradient device as in Truitt *et al.*^14^, we were not able to reproduce the previously obtained results. Keeping the lines in the lab for 4-5 years with high inbreeding prior to the repeated thermal gradient assays might have potentially influenced the results through mutation accumulation, titer reduction or other unknown genetic factors^47^. Moreover, differences in the environmental conditions during our experiment and the experiments in Truitt *et al.*^14^ may strongly confound thermal behavior. For example, the experiments in Truitt *et al.*^14^ were neither controlled for light cues nor for the effects of humidity nor ambient temperature variation. While the experiments in Arnold *et al.*^15^ were carried out in complete darkness, these assays were also neither controlled for humidity variation nor ambient temperature variation. Such environmental factors may have a strong influence on behavior during experimental assays, which could potentially overwhelm subtle variation in thermal preference.

**Table 2.** Statistical analyses of different environmental factors and experimental design influencing *T*_p_ in *Drosophila melanogaster* infected with different *Wolbachia* types. Significant effects of generalized mixed models (GLMM) inferred by analysis of deviance are highlighted in bold.

| **(A) $T_p$ in adults (plate vs tube device)** |  |  |  |
| --- | --- | --- | --- |
| **Effect** | **$\chi^2$** | ***df*** | ***p*-Value** |
| machine | 748.6 | 4 | **< 2.2e-16** |
| infection | 18.9 | 6 | **4.4E-03** |
| machine*infection | 4.6 | 3 | 0.20 |
| **(B) $T_p$ in adults (humidity)** |  |  |  |
| **Effect** | **$\chi^2$** | ***df*** | ***p*-Value** |
| humidity | 266.8 | 4 | **< 2.2e-16** |
| infection | 15.1 | 6 | **2.0E-02** |
| humidity*infection | 2.2 | 3 | 0.52 |
| **(C) $T_p$ in adults (food)** |  |  |  |
| **Effect** | **$\chi^2$** | ***df*** | ***p*-Value** |
| food | 24.1 | 2 | **5.8E-06** |
| infection | 0.2 | 2 | 0.92 |
| food*infection | 0.2 | 1 | 0.68 |
| **(D) $T_p$ in adults (light)** |  |  |  |
| **Effect** | **$\chi^2$** | ***df*** | ***p*-Value** |
| light | 5.7 | 2 | 6.0E-02 |
| infection | 1.5 | 2 | 4.7E-01 |
| light*infection | 0.9 | 1 | 3.4E-01 |

### Humidity, diet and light substantially influence thermal preference of flies independent of Wolbachia infection

The analyses presented above show that the design of the thermal gradient device plays an important role in *T*_p_ measurements. The tubular device consisted of more spare metal parts than the plate device, which negatively affected the temperature conductivity. Accordingly, we found that the effective temperature range along the gradient was much narrower in the tubular compared to the plate device. The minimum temperature in the tube never decreased below 17°C (maximum temperature: 27°C), whereas the plate reached temperatures as low as 13°C (maximum temperature: 27°C). We therefore tested if the flies were limited by the temperature range and were unable to choose preferred temperatures below the minimum temperature in the tube. In this case, flies may cluster at the lower end of the gradient in the tube device, which would result in *T*_p_ distributions skewed towards the lower end, compared to the plate device. However, when comparing the skewness of *T*_p_ distributions in replicate experiments among the two devices we did not find significant differences (Wilcoxon Rank test, *W*=258, *p*=0.128) which suggests that the temperature range within the tube did not limit the behavior of the test flies.

Conversely, cold temperatures as low as 10°C-13°C may cause reduced velocity^48^ which can lead to immobilization^49, 50^ as a physiological response. This may thus result in biased *T*_p_ measurements due to a clustering of immobilized flies that get trapped at the cold end of the gradient^18^. Similar to Hague et al.^16^, we thus repeated testing for the influence of *Wolbachia* on *T*_p_ in flies on the plate device by excluding flies with a *T*_p_ <15°C, to avoid a potential bias from reduced mobility in cold temperatures. The analysis based on this reduced dataset did not indicate a *Wolbachia*-specific influence on host *T*_p_ either (GLMM: p=0.43, *x*^2^=2.76, *df*=3) and thus did not qualitatively differ from results based on the full dataset. This suggests that trapping due to immobilization at the cold end of the gradient is unlikely a confounding factor in our experiments.

We speculate that also other environmental factors associated with the design of the devices may strongly affect thermal behavior. According to Dillon *et al.*^18^, sex, age, humidity, light, circadian rhythms, feeding status and number of individuals tested at once all influence thermal behavior. While similar numbers of flies that were tested simultaneously during each experiment on both devices, the available space for each fly differed dramatically between the two designs. In particular, the narrower space in the plate device may have led to stronger interactions among the flies which could result in pronounced gregarism and clustering due to social interactions at certain regions along the arenas^51^. To further investigate how experimental design influences *T*_p_ measurement, we compared 15 studies on thermal behavior in *D. melanogaster* flies. This qualitative meta-analysis revealed high variation in measured *T*_p_, which apparently depends on the design of the gradient device, the range or the temperature gradient, rearing and experimental conditions, food, age of flies and number of individuals tested (all summarized in **Table S3**).

To better understand the influence of specific environmental stimuli on thermal preference assays, we experimentally investigated how humidity, food and light affect *T*_p_ measurement in the context of varying *Wolbachia* infection status. We observed a highly significant effect of humidity (GLMM: *p*<2.2e-16, *x*^2^=266.93, *df*=4; **Table 2B**; **Figure 3**) on *T*_p_ of adult flies. Higher humidity (60%) resulted in increased thermal preferences, elevated by 1-2°C (**Table S2)**. The median *T*_p_ at 30% humidity ranged from 21.5°C (*w*MelCs and *w*MelPop) to 22°C (*w*-), whereas the median *T*_p_ at 60% humidity ranged from 22.7°C (*w*-) to 23.5°C (*w*Mel, *w*MelCS). A positive correlation between humidity and *T*_p_ was similarly observed already in other insects^52, 53^.

**Figure 3.**
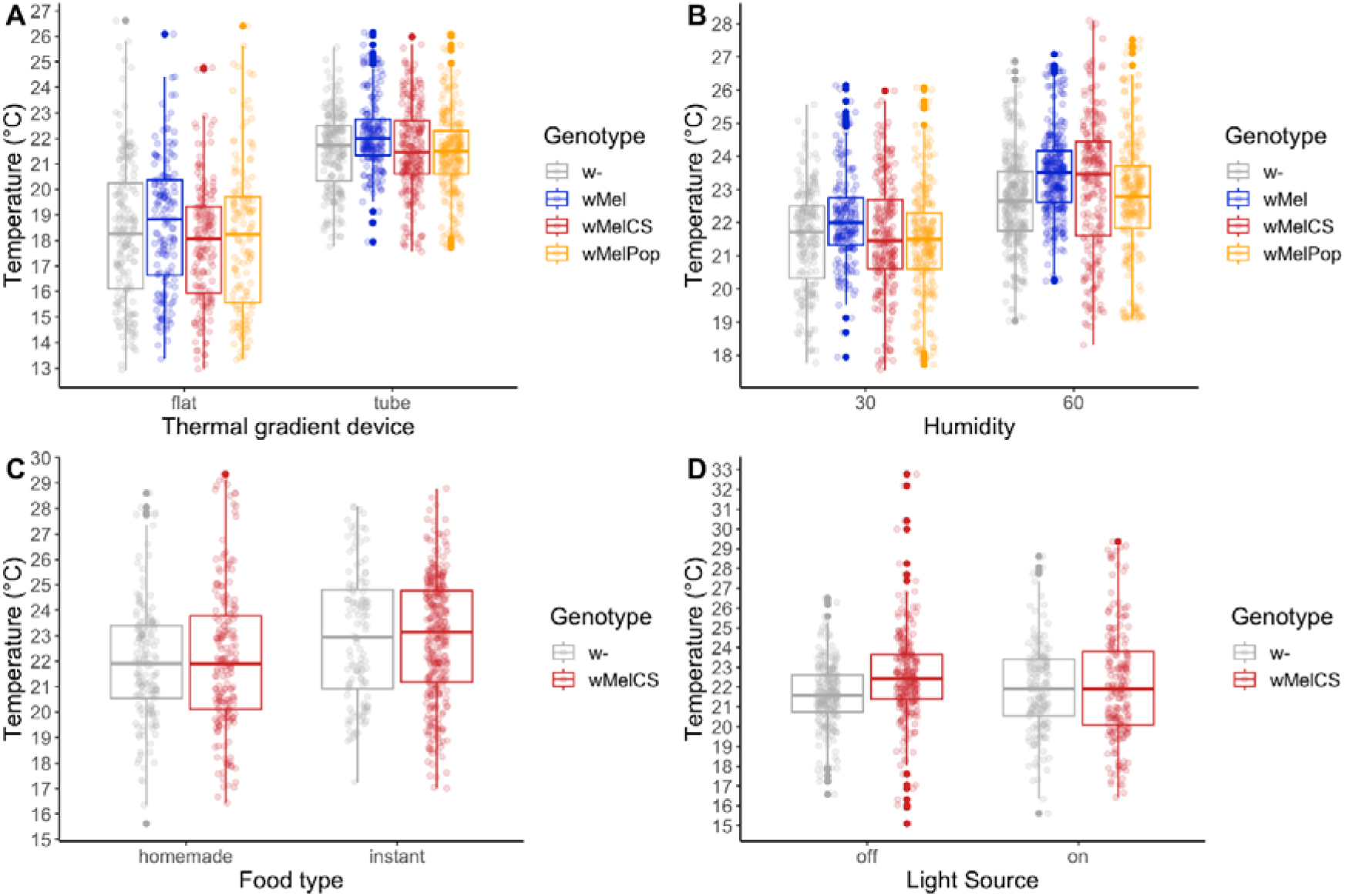
The influence of environmental and experimental factors on *T*_p_ in *Wolbachia*-infected *Drosophila melanogaster*. Boxplots showing the influence of the design of the thermal gradient device, humidity, food, and light on measurement of *T*_p_ in *D. melanogaster* and the influence of *Wolbachia* on host thermal preference, where the colors highlight the different *Wolbachia* genotypes

Diet also had a strong influence on thermal behavior. We found that flies on in-house food preferred 1°C lower temperatures than counterparts maintained on Carolina instant food (GLMM: *p*=5.79e-6, *x*^2^=24.2, *df*=2; **Table 2C**; **Figure 3**). Differences in diet quality or feeding status (fed versus fasting) are known to influence thermal preference in ectotherms, where rich food diet increases *T*_p_ compared to poor food or fasting conditions^54–57^. Our results match these previous findings. While we did not assess the exact nutrient composition in our in-house fly food, the recipe is considered less rich in carbohydrates and proteins than Carolina 4-24 instant food. Additionally, diet is also known to affect the titer of *Wolbachia* in different tissues: Protein-rich food has been found to boost titer levels in somatic tissues, but not in ovaries. Conversely, carbohydrate-rich food can increase titer levels in ovaries only^58, 59^. If variation in titer levels may have a strong influence on thermal preference, as speculated previously^14, 16^, the food quality may indirectly influence *T*_p_ by modulating titer levels. However, we failed to observe a significant influence of light on *T*_p_, (GLMM: *p*=0.059, *x*^2^=5.69, *df*=2; **Table 2D**; **Figure 3**)

Interestingly, in contrast to the experiments performed in our newly designed “plate” gradient device, the follow-up experiments in the “tube”, in particular the experiment testing for the effects of humidity variation, revealed significant effects of *Wolbachia* infections on *T*_p_ (see **Table 2A**). However, the observed patterns were opposite to expectations. Flies infected with *w*MelCS preferred warmer temperatures than uninfected flies in both experiments (Tukey’s post hoc test: *p*<0.01; **Table S2**; **Figure 3**). These findings contradict previously observed results where flies infected with *w*MelCS variant preferred lower temperatures than uninfected counterparts^14, 15^. Our results are, however, in line with the data observed by Hague et al.^16^, showing no shift of preference to higher temperatures in flies artificially infected with *w*MelCS. In summary, these results further suggest that *T*_p_ measurement is strongly influenced by the experimental setup and that *T*_p_ variation with respect to *Wolbachia* infections is probably very subtle.

## Conclusions

In our thermal behavior assays we found that *Wolbachia* does not influence thermal preference in early and late 3^rd^-instar *Drosophila* larvae, consistent with previous studies suggesting that *Wolbachia* has no, or only very weak effects on juvenile traits of *D. melanogaster*. In contrast, we found subtle effects of *Wolbachia* on thermal preference in adult *D. melanogaster* under certain experimental conditions only. These effects were inconsistent with previous data and indicate that *T*_p_ measurement assays are potentially influenced and confounded by uncontrolled environmental cues. Such unwanted behavioral stimuli can overwhelm subtle thermal preference differences during behavioral experiments. As we show in our study, factors such as humidity, diet and design of the gradient device substantially influence measurements of thermal preference in *D. melanogaster* infected with different *Wolbachia* variants. Our findings demonstrate the necessity of a careful experimental design and cautious interpretations of *T*_p_ measurements together with a thorough description of the methods and equipment used to conduct behavioral studies. Our data and findings thus provide important considerations when planning future behavioral assays to assess thermal preference not only in *Drosophila* but also in other insects infected with *Wolbachia*, such as vectors of human diseases like mosquitoes.

## Acknowledgments

We are especially grateful to Wolfgang Miller who helped with the design of the experiment, who provided lab space and who conceptually supported this project. We are also indebted to Marcel Freund at the University of Zürich who not only hand-crafted the thermal gradient devices used in this study but also helped with their design. Moreover, we thank Thomas Flatt, Elisabeth Haring and Roman Arguello for helpful comments on previous versions of this manuscript. We are very grateful to Fabian Gstöttenmayr, Janis Jeschgo, Julia Mras, Elina Koivisto and Jasmin Jester who helped with fly maintenance and fly food cooking. This project was funded by a standalone grant from the Austrian Science Fund (FWF P32275) to Martin Kapun.

## Authors Contributions

Anton Strunov involved in conceptualization, investigation, data curation, visualization, formal analysis, validation, writing—original draft; Charlotte Schoenherr involved in investigation, data curation and writing—review & editing; Martin Kapun involved in conceptualization, formal analysis, visualization, supervision, funding acquisition, writing— review & editing and project administration.

## Data availability

All raw data and the *R* code to carry out the statistical analyses can be found in the Supplementary Information files Strunov_etal_WolbTP_2023_RawData.xlsx and Strunov_etal_WolbTP_2023_Rcode.zip, respectively. A more detailed description of the devices, methods and statistical approaches used in this manuscript can be found online at https://github.com/capoony/DrosophilaThermalGradient

