## Supporting Information for "Wolbachia has subtle effects on thermal preference in highly inbred Drosophila melanogaster which vary with life stage and environmental conditions"

**Supplementary Material**

**Figure S1.** **Thermal preference (*T*_p_) of 120h old *Wolbachia*-infected *D. melanogaster* 3^rd^-instar in extended temperature range** Temperature shift in 3^rd^-instar larvae 120 h after egg laying (AEL) that were either uninfected (w-; grey) or infected with wMelCS (red).

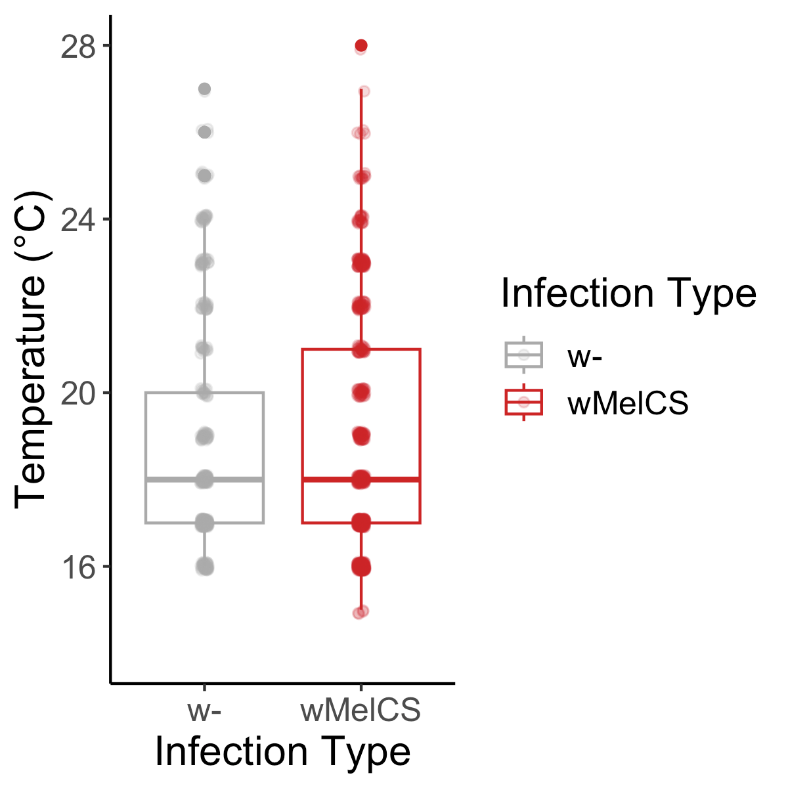

**Table S1**. **Experimental conditions and mean/median *T*_p_ for each factor combination.** Summary table showing the gradient machine design used for the experiments, the number of replicated experiments and the total number of flies used for each type of experiment. Moreover, the table is showing the median and mean *T*_p_ as well as the standard deviation for each of the factor combinations.

| **experiment** | **factor** | **gradient machine used** | **Infection status** | **N of replicates** | **N of individuals** | **Median** | **Mean Tp** | **SD** |
| --- | --- | --- | --- | --- | --- | --- | --- | --- |
| larvae; 72h vs. 120h | 72h | plate (18-30°C) | w- | 3 | 162 | 23 | 22.4 | 2.15 |
|  |  |  | wMelCS | 3 | 119 | 22 | 21.9 | 1.97 |
|  | 120h | plate (18-30°C) | w- | 5 | 364 | 19 | 19.9 | 2.17 |
|  |  |  | wMelCS | 5 | 294 | 19 | 20.2 | 2.59 |
| larvae; 120h | - | plate (15-30°C) | w- | 5 | 332 | 18 | 19 | 2.34 |
|  |  |  | wMelCS | 5 | 529 | 18 | 18.7 | 2.48 |
| larvae; 140h | - | plate (18-30°C) | w- | 5 | 142 | 20 | 20.4 | 1.8 |
|  |  |  | wMelCS | 5 | 192 | 21 | 20.7 | 1.76 |
| adult; plate | - | plate | w- | 5 | 188 | 18.3 | 18.3 | 2.72 |
|  |  |  | wMel | 5 | 157 | 18.8 | 18.7 | 2.37 |
|  |  |  | wMelCS | 5 | 166 | 18.1 | 17.9 | 2.28 |
|  |  |  | wMelPop | 5 | 139 | 18.2 | 18.1 | 2.89 |
| adult; tube | - | tube | w- | 5 | 203 | 21.7 | 21.6 | 1.46 |
|  |  |  | wMel | 5 | 191 | 22 | 22.2 | 1.3 |
|  |  |  | wMelCS | 5 | 235 | 21.5 | 21.6 | 1.67 |
|  |  |  | wMelPop | 5 | 237 | 21.5 | 21.4 | 1.66 |
| adult; humidity | 30% | tube | w- | 5 | 203 | 21.7 | 21.6 | 1.46 |
|  |  |  | wMel | 5 | 191 | 22 | 22.2 | 1.3 |
|  |  |  | wMelCS | 5 | 235 | 21.5 | 21.6 | 1.67 |
|  |  |  | wMelPop | 5 | 237 | 21.5 | 21.4 | 1.66 |
|  | 60% | tube | w- | 3 | 311 | 22.7 | 22.7 | 1.32 |
|  |  |  | wMel | 3 | 293 | 23.5 | 23.5 | 1.24 |
|  |  |  | wMelCS | 3 | 228 | 23.5 | 23.1 | 1.78 |
|  |  |  | wMelPop | 3 | 231 | 22.8 | 22.7 | 1.73 |
| adult; light | on | tube | w- | 3 | 159 | 21.9 | 22 | 2.27 |
|  |  |  | wMelCS | 3 | 193 | 21.9 | 22.1 | 2.34 |
|  | off | tube | w- | 3 | 211 | 21.6 | 21.6 | 1.47 |
|  |  |  | wMelCS | 3 | 243 | 22.4 | 22.3 | 1.99 |
| adult; food | instant | tube | w- | 3 | 159 | 22.9 | 23.1 | 2.47 |
|  |  |  | wMelCS | 3 | 193 | 23.1 | 23 | 2.47 |
|  | homemade | tube | w- | 3 | 121 | 21.9 | 22 | 2.27 |
|  |  |  | wMelCS | 3 | 313 | 21.9 | 22.1 | 2.34 |

**Table S2. Tukey’s post hoc pairwise comparison of experimental or environmental factors that influence the effect of *Wolbachia* on host *T*_p_.** The table is showing significant pairwise comparisons in bold.

| **Test** | **contrast** | **estimate** | **SE** | **df** | **t.ratio** | **p.value** |
| --- | --- | --- | --- | --- | --- | --- |
| Adult; tube | (w-) - (wMel) | -0.583 | 0.221 | 9.91 | -2.643 | 0.0969 |
|  | (w-) - (wMelCS) | -0.0332 | 0.213 | 8.75 | -0.156 | 0.9986 |
|  | (w-) - (wMelPop) | 0.1627 | 0.213 | 8.58 | 0.762 | 0.8691 |
|  | (wMel) - (wMelCS) | 0.5498 | 0.217 | 9.32 | 2.538 | 0.1175 |
|  | **(wMel) - (wMelPop)** | **0.7457** | **0.216** | **9.1** | **3.447** | **0.0302** |
|  | (wMelCS) - (wMelPop) | 0.1959 | 0.21 | 8.05 | 0.934 | 0.7883 |
| Adult; comparison devices | (w-; flat) - (wMel; flat) | -0.188 | 0.297 | 35.5 | -0.633 | 0.9981 |
|  | (w-; flat) - (wMelCS; flat) | 0.6119 | 0.261 | 34.5 | 2.341 | 0.3016 |
|  | (w-; flat) - (wMelPop; flat) | 0.1814 | 0.283 | 39.1 | 0.641 | 0.998 |
|  | **(w-; flat) - (w-; tube)** | **-3.1768** | **0.227** | **975.2** | **-13.996** | **<.0001** |
|  | **(w-; flat) - (wMel; tube)** | **-3.7717** | **0.258** | **30.2** | **-14.64** | **<.0001** |
|  | **(w-; flat) - (wMelCS; tube)** | **-3.2149** | **0.251** | **26.4** | **-12.798** | **<.0001** |
|  | **(w-; flat) - (wMelPop; tube)** | **-3.0293** | **0.243** | **25.9** | **-12.475** | **<.0001** |
|  | (wMel; flat) - (wMelCS; flat) | 0.7999 | 0.286 | 38.2 | 2.793 | 0.1268 |
|  | (wMel; flat) - (wMelPop; flat) | 0.3694 | 0.281 | 43 | 1.314 | 0.8886 |
|  | **(wMel; flat) - (w-; tube)** | **-2.9888** | **0.268** | **32.6** | **-11.155** | **<.0001** |
|  | **(wMel; flat) - (wMel; tube)** | **-3.5837** | **0.238** | **1115.3** | **-15.034** | **<.0001** |
|  | **(wMel; flat) - (wMelCS; tube)** | **-3.027** | **0.253** | **29.3** | **-11.944** | **<.0001** |
|  | **(wMel; flat) - (wMelPop; tube)** | **-2.8413** | **0.261** | **28.2** | **-10.88** | **<.0001** |
|  | (wMelCS; flat) - (wMelPop; flat) | -0.4305 | 0.296 | 42.6 | -1.457 | 0.8253 |
|  | **(wMelCS; flat) - (w-; tube)** | **-3.7887** | **0.263** | **31.2** | **-14.388** | **<.0001** |
|  | **(wMelCS; flat) - (wMel; tube)** | **-4.3836** | **0.258** | **32.7** | **-16.971** | **<.0001** |
|  | **(wMelCS; flat) - (wMelCS; tube)** | **-3.8269** | **0.236** | **521.1** | **-16.185** | **<.0001** |
|  | **(wMelCS; flat) - (wMelPop; tube)** | **-3.6412** | **0.252** | **28.7** | **-14.448** | **<.0001** |
|  | **(wMelPop; flat) - (w-; tube)** | **-3.3582** | **0.26** | **35.6** | **-12.913** | **<.0001** |
|  | **(wMelPop; flat) - (wMel; tube)** | **-3.9531** | **0.267** | **37.9** | **-14.825** | **<.0001** |
|  | **(wMelPop; flat) - (wMelCS; tube)** | **-3.3964** | **0.252** | **32.6** | **-13.464** | **<.0001** |
|  | **(wMelPop; flat) - (wMelPop; tube)** | **-3.2107** | **0.236** | **1100.8** | **-13.587** | **<.0001** |
|  | (w-; tube) - (wMel; tube) | -0.5949 | 0.242 | 26.7 | -2.455 | 0.2571 |
|  | (w-; tube) - (wMelCS; tube) | -0.0381 | 0.232 | 22.8 | -0.165 | 1 |
|  | (w-; tube) - (wMelPop; tube) | 0.1475 | 0.231 | 22.1 | 0.638 | 0.9978 |
|  | (wMel; tube) - (wMelCS; tube) | 0.5567 | 0.236 | 24.5 | 2.363 | 0.3019 |
|  | (wMel; tube) - (wMelPop; tube) | 0.7424 | 0.235 | 23.7 | 3.157 | 0.07 |
|  | (wMelCS; tube) - (wMelPop; tube) | 0.1857 | 0.226 | 19.9 | 0.821 | 0.9897 |
| Adult; Food | (w-; homemade) - (wMelCS; homemade) | -0.0956 | 0.42 | 3.16 | -0.228 | 0.995 |
|  | **(w-; homemade) - (w-; instant)** | **-1.002** | **0.3** | **777.08** | **-3.34** | **0.0049** |
|  | (w-; homemade) - (wMelCS; instant) | -0.9481 | 0.407 | 2.77 | -2.327 | 0.2817 |
|  | (wMelCS; homemade) - (w-; instant) | -0.9064 | 0.435 | 3.61 | -2.086 | 0.3075 |
|  | **(wMelCS; homemade) - (wMelCS; instant)** | **-0.8525** | **0.233** | **779.84** | **-3.667** | **0.0015** |
|  | (w-; instant) - (wMelCS; instant) | 0.0539 | 0.422 | 3.19 | 0.128 | 0.9991 |
| Adult; Humidity | (w-; 30) - (wMel; 30) | -0.5198 | 0.217 | 19.7 | -2.394 | 0.2966 |
|  | (w-; 30) - (wMelCS; 30) | -0.0106 | 0.211 | 17.5 | -0.05 | 1 |
|  | (w-; 30) - (wMelPop; 30) | 0.166 | 0.212 | 17.5 | 0.784 | 0.9919 |
|  | **(w-; 30) - (w-; 60)** | **-1.3024** | **0.156** | **1151.3** | **-8.355** | **<.0001** |
|  | **(w-; 30) - (wMel; 60)** | **-1.9635** | **0.222** | **15.6** | **-8.855** | **<.0001** |
|  | **(w-; 30) - (wMelCS; 60)** | **-1.5737** | **0.225** | **18.3** | **-6.993** | **<.0001** |
|  | **(w-; 30) - (wMelPop; 60)** | **-1.1208** | **0.222** | **18.4** | **-5.054** | **0.0016** |
|  | (wMel; 30) - (wMelCS; 30) | 0.5092 | 0.213 | 18.4 | 2.392 | 0.3007 |
|  | (wMel; 30) - (wMelPop; 30) | 0.6858 | 0.214 | 18.4 | 3.21 | 0.0727 |
|  | **(wMel; 30) - (w-; 60)** | **-0.7825** | **0.221** | **17.4** | **-3.539** | **0.0401** |
|  | **(wMel; 30) - (wMel; 60)** | **-1.4437** | **0.167** | **554** | **-8.633** | **<.0001** |
|  | **(wMel; 30) - (wMelCS; 60)** | **-1.0539** | **0.225** | **18.8** | **-4.674** | **0.0034** |
|  | (wMel; 30) - (wMelPop; 60) | -0.601 | 0.223 | 18.9 | -2.696 | 0.1836 |
|  | (wMelCS; 30) - (wMelPop; 30) | 0.1766 | 0.208 | 16.2 | 0.85 | 0.9868 |
|  | **(wMelCS; 30) - (w-; 60)** | **-1.2918** | **0.214** | **15.4** | **-6.023** | **0.0004** |
|  | **(wMelCS; 30) - (wMel; 60)** | **-1.953** | **0.215** | **15** | **-9.071** | **<.0001** |
|  | **(wMelCS; 30) - (wMelCS; 60)** | **-1.5631** | **0.172** | **644.7** | **-9.114** | **<.0001** |
|  | **(wMelCS; 30) - (wMelPop; 60)** | **-1.1102** | **0.218** | **17.1** | **-5.101** | **0.0018** |
|  | **(wMelPop; 30) - (w-; 60)** | **-1.4684** | **0.216** | **15.8** | **-6.809** | **0.0001** |
|  | **(wMelPop; 30) - (wMel; 60)** | **-2.1295** | **0.223** | **15.5** | **-9.558** | **<.0001** |
|  | **(wMelPop; 30) - (wMelCS; 60)** | **-1.7397** | **0.222** | **18.8** | **-7.826** | **<.0001** |
|  | **(wMelPop; 30) - (wMelPop; 60)** | **-1.2868** | **0.172** | **647.4** | **-7.496** | **<.0001** |
|  | (w-; 60) - (wMel; 60) | -0.6612 | 0.224 | 13 | -2.955 | 0.1387 |
|  | (w-; 60) - (wMelCS; 60) | -0.2713 | 0.226 | 14.9 | -1.201 | 0.9196 |
|  | (w-; 60) - (wMelPop; 60) | 0.1816 | 0.222 | 14.6 | 0.817 | 0.9892 |
|  | (wMel; 60) - (wMelCS; 60) | 0.3898 | 0.233 | 14 | 1.671 | 0.704 |
|  | **(wMel; 60) - (wMelPop; 60)** | **0.8428** | **0.227** | **14** | **3.709** | **0.0362** |
|  | (wMelCS; 60) - (wMelPop; 60) | 0.4529 | 0.229 | 17.1 | 1.978 | 0.5223 |

**Table S3. Meta-analysis of studies investigating *T*_p_ in *Drosophila*.** Observations of *T*_p_ from literature with detailed information about physical parameters and environmental conditions used in a study.

|  | **fly strain** | **observed Tp** | **Tp device** | **gradient range** | **rearing  conditions** | **experiment conditions** | **food** | **age of flies** | **N of flies  per test group** | **source** |
| --- | --- | --- | --- | --- | --- | --- | --- | --- | --- | --- |
| **1** | D. melanogaster  w1118 (generated by Luis Teixeira) | 24.4°C | 60 cm long polycarbonate tube from Rajpurohit and Schmidt, 2016 | 16 - 36°C | 25°C 45% humidity 12h light/dark | 24°C RT 40% humidity in a light room | *Drosophila* Formula 4‐24®  Instant Medium (Carolina®, NC) | 3 days, 1 week, 2 weeks no age effect | 75-100 mixed sex | Truitt et al., 2018 Environmental microbiology |
| **2** | D. melanogaster  w1118 | 25°C | aluminium plate 12.5x20 cm | 13 - 33°C | 25°C 12h light/dark | 17°C RT 70% humidity in a light rom | standard cornmeal diet | 0-3 days | 30 mixed sex | Ueno et al., 2012 PloS One |
| **3** | D. melanogaster  w1118 | peak at 26°C | aluminium plate 50x30 cm 4 lanes | 10 - 40°C | 25°C 70% humidity 12h light/dark | 24-26°C RT 40-50% humidity in darkness | standard cornmeal diet | different age Tp is age dependent | 80-120 mixed sex | Shih et al., 2015 |
| **4** | D. melanogaster  w1118 | 25°C in light 24°C in dark | aluminium slab from Sayeed and Benzer, 1996 | 18 - 32°C | 25°C 12h light/dark | 25°C RT 65-70% humidity light/dark | standard cornmeal diet | 0-3 days | 20-30  mixed sex | Head et al., 2015 Current biology |
| **5** | D. melanogaster  w1118 | 24-25°C | aluminium slab 40 cm | 15-45°C | 24-25°C 12h light/dark | 25°C RT 40-50% humidity in darkness | standard cornmeal diet | 5-5 days | 45-70 mixed sex | Hong et al., 2006 J Neuroscience |
| **6** | D. melanogaster  Canton S | 1) peak at 24°C 2) peak at 23°C | aluminium slab 27 cm 4 lanes | 1) 18 - 35°C 2) 23 - 36.5°C | 25°C 60% humidity 12h light/dark | in darkness moist Whatman paper | standard cornmeal diet | ND | 10 flies per lane mixed sex | Sayeed and Benzer, 1996 PNAS |
| **7** | D. melanogaster  Canton S | 22.5°C | aluminium slab 50 cm 5 lanes | 14 - 30°C | 25°C 12h light/dark | 27% humidity | standard cornmeal diet | ND | 1 fly per lane | Giraldo et al., 2019 Scientific reports |
| **8** | D. melanogaster  Oregon RC (treated with 0.03% tetracyclne and restored) | 25.8°C ± 0.24 | polycarbonate tube from Arnold et al., 2015 | 17.5 - 33.5°C | 25°C humidity - ND 12h light/dark | in darkness between 9.30  and 13.30 | standard cornmeal diet | 4-7 days | 5  only males | Arnold et al., 2019 Ecological Entomology |
| **9** | D. melanogaster | 25°C | aluminium slab from Sayeed and Benzer, 1996 | 18 - 32°C | ND | 25°C RT 70% humidity in darkness | ND | 0-3 days | 20-30  mixed sex | Hamada et al., 2008 Nature |
| **10** | D. melanogaster originally from Denmark | 26°C | aluminium slab 156 cm 8 lanes | 9 - 34°C | 19°C 12h light/dark | ND | standard cornmeal diet | 1-4 days | 60 mixed sex | MacLean et al., 2019 J Therm Biology |
| **11** | D. melanogaster Orlando, USA wild-caught outbred stock | 26.3°C | aluminium slab 90 cm | 18 - 32°C | 25°C 12h light/dark | ND | standard cornmeal diet | 5 days | 1 virgin females | Fedorka et al., 2016 J Insect Physiology |
| **12** | D. melanogaster wild-caught Japan | 21.8°C | glass tube 78 cm | 5-35°C | 25°C | ND | standard cornmeal diet | different age | 20 separate sex | Yamamoto and Ohba, 1984 Zool Sci |
| **13** | D. melanogaster wild-caught Cambridge isofemale line | 23.1°C | aluminium slab 16 lanes | 18-30°C | 25°C 30-40% humidity 12h light/dark | in a light room | standard cornmeal diet | ND | 1 fly per lane mixed sex | Kain et al., 2015 Evolution |
| **14** | D. melanogaster wild-caught east coast of USA | 20-21°C | 60 cm long polycarbonate tube | 12 - 32°C | 25°C 12h light/dark | ND | standard cornmeal diet | 4-5 days | 150 each sex separate | Rajpurohit and Schmidt, 2010 Fly |
| **15** | D. melanogaster wild-caught with wMel and labstrain with wMelCS | 25°C for wMel 27°C for wMelCS | 10 cm long lane on a platform | 17 - 34°C | 25°C 12h light/dark | cold room, 5°C | standard cornmeal diet | 3-5 days | 50-60 each sex separate (virgins!) | Hague et al., 2020, mBio |

**Supp. raw data table.** Strunov_etal_WolbTP_2023_RawData.xlsx
